# Decoding familiar visual object categories in the mu rhythm oscillatory response

**DOI:** 10.1101/2023.07.17.548986

**Authors:** Kerri M Bailey, Saber Sami, Fraser W Smith

## Abstract

Whilst previous research has linked attenuation of the mu rhythm to the observation of specific visual categories, and even to a potential role in action observation via a putative mirror neuron system, much of this work has not considered what specific type of information might be coded in this oscillatory response when triggered via vision. Here, we sought to determine whether the mu rhythm contains content-specific information about the identity of familiar (and also unfamiliar) graspable objects. In the present study, right-handed participants (*N*=27) viewed images of both familiar (apple, wine glass) and unfamiliar (cubie, smoothie) graspable objects, whilst performing an orthogonal task at fixation. Multivariate pattern analysis (MVPA) revealed significant decoding of familiar, but not unfamiliar, visual object categories in the mu rhythm response. Thus, simply viewing familiar graspable objects may automatically trigger activation of associated tactile and/or motor properties in sensorimotor areas, reflected in the mu rhythm. In addition, we report significant attenuation in the central beta band for both familiar and unfamiliar visual objects, but not in the mu rhythm. Our findings highlight how analysing two different aspects of the oscillatory response – either attenuation or the representation of information content – provide complementary views on the role of the mu rhythm in response to viewing graspable object categories.

**Highlights:**

- The Mu Rhythm oscillation contains fine-grained information about the identity of familiar, graspable objects (but not unfamiliar)
- This study offers evidence of a possible oscillatory marker for cross-sensory effects involving sensorimotor and visual cortices
- Different analysis techniques (univariate vs multivariate) imply different conclusions about the role of specific oscillations in the perception of graspable objects
- The alpha rhythm at occipital sites shows greater attenuation for unfamiliar objects but better representation for familiar objects consistent with sharpening accounts of Predictive Processing

## Introduction

The mu rhythm is a neural oscillation measured over sensorimotor areas that desynchronizes when participants are engaged in executing an action, observing an action, or even when one merely has the intention to act (Fox et al., 2016; Muthukumaraswamy and Johnson, 2004; Pfurtscheller et al., 1997; Pineda, 2005). As such, the mu rhythm has often been considered to be an index of mirror neuron activity (Fox et al., 2016; Muthukumaraswamy et al., 2004). However, recently this has become a problematic interpretation (Coll et al., 2017; Hobson and Bishop, 2016). Importantly, the mu rhythm shows greater desynchronization when participants view pictures of tools vs non-tools (Proverbio, 2012) and when participants view real 3D objects vs pictures (Marini et al., 2019). In both these cases greater action affordances (Gibson, 1979; Warman 2021) may be argued to drive the larger mu response to viewing objects. Despite this, it is unclear from these studies what information content is represented in the mu rhythm (i.e. how specific the response is to different visual categories) and whether the response depends upon familiarity with the objects presented. At present it is possible that the larger mu responses for one category vs another (e.g. tools vs non-tools) merely reflect high level effects of attention or arousal to different categories of stimuli. If the mu rhythm carries content specific information relevant to particular objects (and associated actions or tactile properties), i.e. which would be useful for future interaction with such objects, then one should be able to discriminate between such objects based on its response (see Coll et al., 2017). In the present experiment, for the first time, we explicitly test the information contained in the mu rhythm while different graspable object categories are presented to participants. Moreover, in the above cited studies the stimuli in both object categories were familiar to the participants. Thus, it is unclear whether familiarity is required to elicit an effect on the mu rhythm when triggered via vision or whether one might find similar effects even for unfamiliar but graspable objects.

While the mu rhythm is most often associated with action (either executed or observed) in fact there are several reports that it may instead be more related to tactile stimulation than action per se (Cheyne et al., 2003; Coll et al., 2017; Ritter et al., 2009). For example, research by Cheyne et al. (2003) used MEG to find tactile stimulation during a finger brushing task produced suppression of the mu rhythm. In our recent work using fMRI, we have shown that viewing familiar but not unfamiliar graspable objects leads to content-specific information being present in primary somatosensory areas (Smith and Goodale, 2015; see also Bailey et al., 2023), the first cortical areas dedicated to processing our sense of touch. At present however, it remains unclear whether the mu response (either its desynchronization or the information content) is sensitive to familiarity with the presented objects. To the extent that the mu response is generated within SI (as opposed to M1) we would expect to able to detect a similar effect as we have found in fMRI.

It is also worth noting other oscillations of interest in the present study, such as the central beta and occipital alpha oscillations. First, the functional roles of the mu rhythm in comparison to the central beta oscillation are not well understood (Cheyne, 2013). Both oscillations have been suggested to originate from deep layers of cortex (Bastos et al., 2015; Michalareas et al., 2016), thus both rhythms may actually reflect aspects of cortical feedback processing rather than bottom-up signalling. However, one interesting difference between the mu rhythm and the central beta oscillation was found in work by Cheyne (2013), who suggested the mu rhythm reflects activity in SI, while the central beta rhythm reflects activity in primary motor cortex (MI). As such, we may only expect to find discriminable information within the mu rhythm, since this is the oscillation which has been suggested to best reflect underlying activity in SI – especially since Smith and Goodale (2015) did not find discriminable patterns of information in M1 when participants viewed the same familiar visual objects. Second, the occipital alpha oscillation is known to reflect neuronal top-down influences on perception (Sherman et al., 2016; Xie et al 2020; Chen et al 2023), therefore investigating differences in this frequency band in relation to prior experience with graspable objects is also of interest. In the predictive coding framework (Clark 2013) alpha oscillations are thought to convey predictions sent back to earlier sensory areas (Bastos et al 2012) and as such we may expect information contained within this band over occipital sites to be especially sensitive to effects of prior knowledge – i.e. familiarity.

In the present study, we used multivariate pattern analysis (MVPA) to determine whether the mu rhythm contains content-specific information about the identity of viewed familiar and unfamiliar graspable objects. We investigated familiarity with objects to determine whether experience with the haptic interactions of the object is necessary to observe such effects in the mu rhythm. Since links have already been found between vision and SI when using fMRI (Meyer and Damasio, 2009; Smith and Goodale, 2015), and given that recently the mu rhythm has been proposed to reflect tactile aspects of object interaction or tactile stimulation (e.g. Cheyne, 2013; Coll et al., 2017), we expect to find a similar effect can be detected in the mu rhythm using EEG. We also investigated whether such effects could be detected in the beta (15-25 Hz) frequency band (as in Coll et al., 2017). The main analysis technique employed was MVPA, since it indicates the information content from the task (Mur et al., 2009), as opposed to univariate analysis which can only investigate overall involvement in a task based on changes in synchronisation in a given region. We expected to find discriminable patterns of information related to the different familiar, but not unfamiliar, visual object categories in the mu rhythm oscillatory response.

## Methods

### Participants

Participants (*N* = 27; 13 male) were right handed (Oldfield, 1971), with an age range of 18-34 years (*M* = 21.19, *SD* = 3.35). All participants reported normal or corrected-to-normal vision and no history of neurological disorders. Written consent was obtained in accordance with approval from the Research Ethics Committee of the School of Psychology at the University of East Anglia and in accord. Participants were recruited through an online system and awarded partial course credit, or through a paid participant panel, receiving either 8 course credits or £16 respectively for their participation.

### Stimuli and design

Two different conditions of full colour visual object stimuli were used in a block design: familiar or unfamiliar graspable objects. Familiar objects consisted of apples and wine glasses (Smith & Goodale, 2015; see Figures 1A and 1B), and unfamiliar objects consisted of cubies and smoothies (Op de Beeck, Torfs, & Wagemans, 2008; see Figures 1C and 1D). There were three exemplars of each visual object, resulting in 12 individual stimuli total. Familiar objects were chosen based on the assumption that participants would have a rich haptic experience with such objects (as in Smith and Goodale, 2015). All images were 400 x 400 pixels, displayed against a white background in the centre of a 24” monitor screen (resolution 1920 x 1080 pixels) using E-Prime 2.0. A black and white fixation cross with a black border (28 x 28 pixels), and a red and green fixation cross with a black border (28 x 28 pixels), was also used. A viewing distance of 45cm was maintained for a visual angle height of 14° for the stimuli, as in Smith and Goodale (2015).

**Figure 1:**
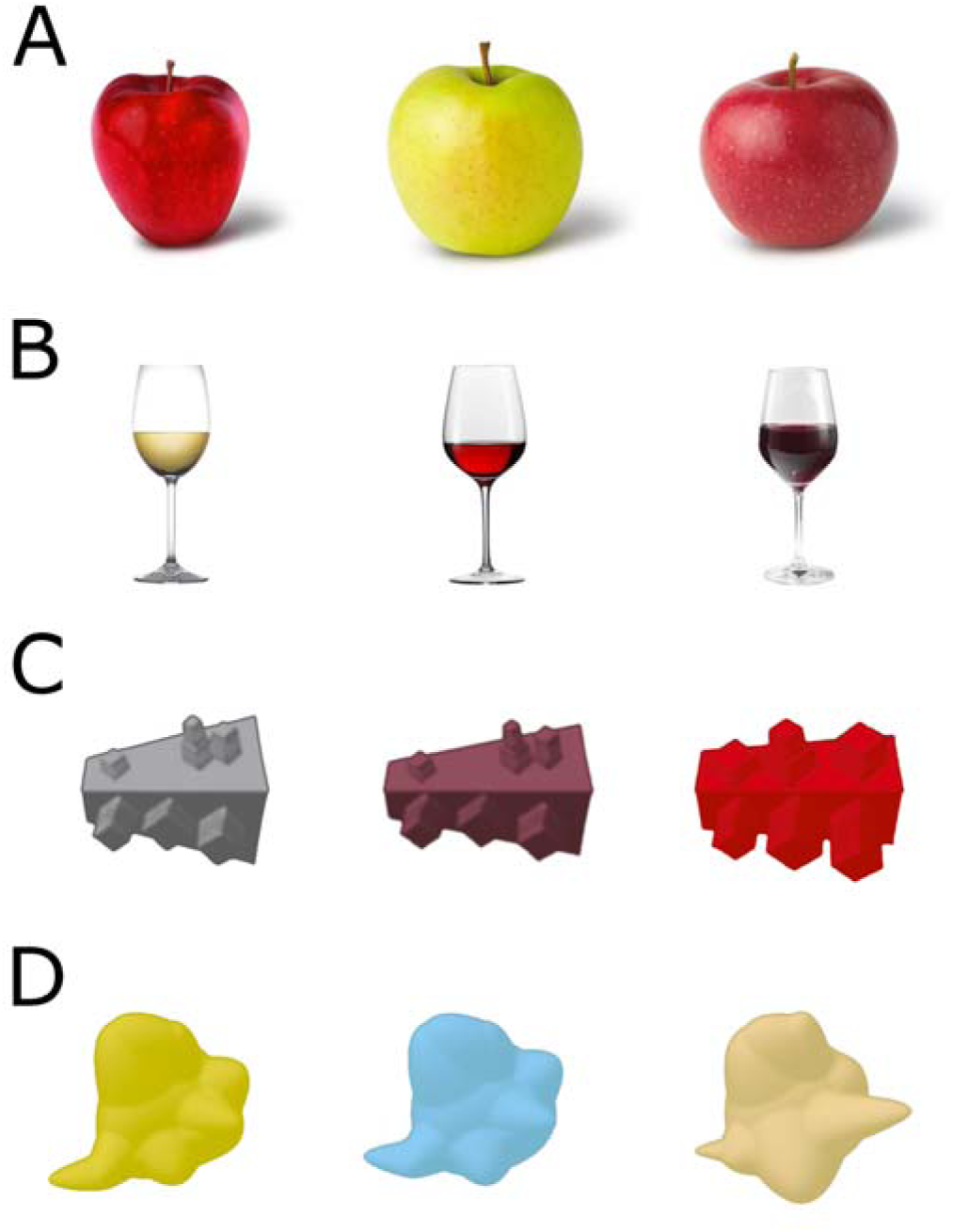
Familiar visual object categories; (A) three exemplars of an apple, (B) three exemplars of a wine glass. Unfamiliar visual object categories; (C) three exemplars of a cubie, (D) three exemplars of a smoothie. Familiar objects taken from Smith and Goodale (2015), and Unfamiliar objects modified from Op de Beeck, Torfs, and Wagemans (2008).

### Apparatus and materials

A 64-channel slim active electrode system (Brain Products GmbH: BrainVision actiCAP) with a BrainAmp MR64 PLUS amplifier was used for the EEG data acquisition (see EEG data acquisition below for more information). The Karolinska Sleepiness Scale (KSS; Akerstedt, Anund, Axelsson, & Kecklund, 2014; Åkerstedt & Gillberg, 1990) was used to measure participant’s sleepiness during the five minutes prior to the end of the experiment. This scale is rated on a Likert scale from 1-9 with the following options: 1) extremely alert, 2) very alert, 3) alert, 4) rather alert, 5) neither alert nor sleepy, 6) some signs of sleepiness, 7) sleepy, but no effort to keep awake, 8) sleepy, some effort to keep awake, and 9) very sleepy, great effort to keep awake, fighting sleep. We included this scale to test whether participants attention and/or drowsiness correlated with overall levels of occipital alpha activity (Niedermeyer and da Silva, 2005).

### Procedure

Upon arrival, participants signed informed consent and the EEG cap was mounted. Once the cap was installed, participants received both verbal and written instructions for the task and were trained via practice trials. Before beginning the experiment, each participant was shown their eye blinks and muscle artifacts (e.g. teeth grinding) on the EEG monitor to emphasize the importance of remaining still during EEG recording, and was asked to blink between trials where possible. During the experiment, participants sat in a dimly lit room. A black and white fixation cross with a black border remained in the centre of the screen throughout the entirety of a block, and each block began and ended with 2000ms of fixation against a white background. After the initial 2000ms fixation, 12 individual stimuli were displayed in a randomly allocated order, meaning each unique stimulus was presented once per block. Each stimulus trial began with 1000ms of fixation, followed by a stimulus presentation of 1000ms whereby the fixation remained on the screen. Each stimulus offset followed a variable delay of fixation for 1500-1900ms. There were 50 blocks in total, however 10 of which were catch blocks. In a catch block, a red and green fixation cross with a black border (28 x 28 pixels) was displayed over one of the stimuli at random.

Participants were instructed to remain fixated on the central fixation cross in order to detect whether there was a colour change in the fixation cross during a block. Participants were asked to pay attention to the stimuli which would appear behind the fixation cross, but to remain fixated at all times. At the end of an experimental block, a question screen appeared which asked participants whether they had detected a red and green fixation cross. Participants were instructed to press one of two buttons on a four-button response device with their right hand in order to answer yes or no, thus eliminating the need for a motor response during experimental trials (see also Smith & Goodale, 2015 for a similar approach). Participants would receive a break screen after their response, enabling them to take a break before the next block if they wished to do so. Every 10 blocks, participants received a longer break in which the screen offset was controlled by the experimenter. During this break, the participant was checked on by the experimenter and offered water to aid alertness during the experiment. In total, each participant was exposed to 40 repetitions of each unique stimulus, resulting in 240 familiar visual objects, and 240 unfamiliar visual objects after removing the catch blocks. The main experiment lasted approximately 40-50 minutes.

At the end of the main experiment, participants were asked to complete the KSS (see Apparatus and materials above) to indicate their sleepiness on a scale from 1 (extremely alert) to 9 (very sleepy, great effort to keep awake, fighting sleep) during the five minutes before completing the rating. This was included since attention and/or drowsiness has previously been found to correlate with overall levels of occipital alpha activity (Niedermeyer and da Silva, 2005). Then, since a key feature of mu suppression is its occurrence both when an individual observes or executes an action (Pfurtscheller et al., 1996), participants were asked to complete a voluntary motor response task in order to map the mu rhythm in relation to a physical motor response. In this experiment, a central black and white fixation cross with a black border remained in the centre of the screen with a white background throughout the entire block. Participants were instructed to fixate on the black and white central fixation cross and press one button from a four-button response device with their right index finger approximately every six seconds. The fixation cross was included to ensure task engagement when making button responses. Each participant was encouraged not to count in their head to avoid alpha contamination (Hobson and Bishop, 2016). Participants completed 40 trials of the button pressing, separated in to four blocks of 10 trials which lasted approximately 5-10 minutes total. On completion of the voluntary motor response task, the EEG cap was dismounted and hair washing facilities were offered to all participants. Participants were debriefed before leaving the room. The entire session lasted no more than two hours per participant.

### EEG data acquisition

The electroencephalogram (EEG) was recorded with a 64-channel slim active electrode system (Brain Products GmbH: BrainVision actiCAP) embedded in a nylon cap (international 10/20 localisation system), with a BrainAmp MR64 PLUS amplifier. One electrode (FT9) was removed from the cap and placed diagonally below and away from the outer canthus of the left eye in order to monitor vertical eye movements (lower EOG). The reference electrode was placed on the tip of the participant’s nose (as in Pfurtscheller, Neuper, Andrew, & Edlinger, 1997), and electrode FT10 was moved to the location of FCz, in order to record from the position where the reference electrode is usually embedded in this cap. The ground electrode was located in the position of FPz. The continuous EEG signal was acquired at a high sampling rate of 1000 Hz. Impedance was kept equal to or less than 50kΩ before recording started.

### EEG data pre-processing

All EEG data pre-processing was performed using the open toolbox EEGLAB (Delorme and Makeig, 2004), used within MATLAB (The MathWorks, USA, 2017b). Raw data from both the main experiment and voluntary motor response task were pre-processed according to the following steps. First, imported data were and high- and low-pass filtered between 0.1 Hz and 50 Hz to remove low-frequency drifts and high-frequency noise respectively. All practice trials, catch blocks, and break periods were then manually removed from the continuous data. A 50 Hz notch filter was also applied to reduce electrical noise. A vertical EOG (VEOG) was reconstructed offline by subtracting the lower EOG from FP1 activity. A horizontal EOG (HEOG) was also constructed by subtracting F7 from F8 activity (Renoult et al., 2015). Independent component analysis (ICA) was then ran on the data to detect eye blink artifacts, which were clearly identified and removed when inspecting components and scalp maps. Finally, data was cleaned using the artifact subspace reconstruction (ASR) plugin developed by Kothe and Jung (2016; Patent No. 14/895,440). See also Chang, Hsu, Pion-Tonachini, and Jung (2018) and Mullen et al. (2013). The ASR interpolated artifact bursts with a variance of more than 5 standard deviations different from the automatically detected clean data (see Gabard-Durnam, Mendez Leal, Wilkinson, & Levin, 2018; Grummett et al., 2014). Data segments post-ASR were then rejected with a time-window rejection criterion of 0.25, meaning the segment was rejected if more than 0.25 of channels were marked as bad. Any channels marked as bad in this process were interpolated, and any remaining noisy electrodes were interpolated based on computing kurtosis with a threshold of 5 using spherical method (Bigdely-Shamlo et al., 2015). This resulted in an average of 2.59 electrodes interpolated (*SD* = 1.72, range 0-5). The cleaned data was then epoched from −1000ms to 1500ms, time-locked to stimulus onset (or time-locked to the button press if the voluntary motor response task), with a baseline correction of −1000ms. Finally, step-like artifact detection was carried out on all electrodes on the epoched data (with the exception of the lower EOG electrode) using a threshold of 100μV and moving window of 200ms in 50ms steps (see Luck, 2005). An average of 0.39% (*SD* = .01, range 0 – 5.21%) of trials were rejected during the entire cleaning process.

### Regions of interest

Two regions of interest (ROI)’s were created for both the univariate and multivariate pattern analysis (MVPA). In line with Coll et al. (2017), ten central electrodes were selected for the central ROI (C1-2-3-4-z, CP1-2-3-4-z). Furthermore, eight occipital electrodes were selected for the occipital ROI (PO3-4-7-8-z, O1-2-z). The central ROI was created to compare the effects of mu rhythm suppression and content specificity to the different visual object categories in each condition (familiar, or unfamiliar visual objects). The occipital ROI was created as a control region to rule out potential confounds by changes in attentional engagement from occipital alpha activity (see Hobson & Bishop, 2016).

### Data analysis

#### Behavioural analysis

Behavioural data for the main experiment was analysed based on correct detection of a red and green fixation cross, or correct rejection of no colour change during a block. Any failures to detect a colour change, or detection of an absent red and green fixation cross was classified as an incorrect response. An average accuracy was calculated for each participant.

#### Univariate cluster-based analysis

Univariate time-frequency analysis was conducted at the channel level by computing the power of event-related desynchronization/synchronization (ERD/ERS) using event-related spectral perturbations (ERSP)’s. The ERSP is a known measure of average dynamic changes in amplitude of the specified broad band EEG frequency spectrum, as a function of time relative to an experimental event (Cuellar and Toro, 2017; Makeig, 1993; Pfurtscheller and Lopes, 1999). To obtain the time-frequency data, a short-time Fourier transform was computed across the averaged baseline-corrected trials by extracting 200 time points in steps of 10ms, using a Hanning-tapered sliding time window with a fixed length of 500ms, covering the entire epoch from −1000ms to 1500ms. Here, a divisive baseline was used relative to the −1000ms to 0ms time period (Ciuparu and Mureşan, 2016; Marini et al., 2019). Power was calculated from 1-30 Hz in steps of exactly 1 Hz. Such analyses were conducted in both central and occipital ROIs and applied to both conditions of familiar and unfamiliar visual object categories, in addition to the voluntary motor response task. Mean changes in spectral power are expressed in decibels (dB).

To avoid circular inference (see Kriegeskorte, Lindquist, Nichols, Poldrack, & Vul, 2010; Kriegeskorte, Simmons, Bellgowan, & Baker, 2009), oscillation clusters were then identified by selecting all pixels across the time-frequency plots during stimulus duration (0-30 Hz, 100 time points corresponding to 0-1000ms) in both the central and occipital ROIs which were statistically significant based on the average ERSP data of *all conditions* (Cohen, 2014), at a significance level of 0.01. Once the precise boundaries of the significant clusters had been defined, the cluster masks were applied separately to each condition (familiar and unfamiliar visual objects) and ROI (central and occipital). The data from each mask and each participant was then extracted and averaged. The average ERSP data was also extracted from the voluntary motor response task by identifying significant clusters based on the ERSP data from the button press.

Overall, a single averaged ERSP value was extracted in each ROI and condition from the mask created from each significant cluster. Paired-sample parametric *t*-tests were then conducted to compare differences between ERSP data for familiar and unfamiliar visual objects. A single ERSP value was also extracted from each cluster and ROI from the voluntary motor response task for each participant. Effect sizes for all *t*-tests are calculated as Cohen’s *d* = *t* / √ N (Lakens, 2013).

#### Univariate time-frequency window analysis

The univariate ERSP data from the main experiment (see Univariate cluster-based analysis above for information on how this data was extracted) was also analysed for significant desynchronization within strictly selected alpha and beta frequency bands for each ROI (central and occipital) averaged over stimulus duration (0-1000ms time-locked to stimulus onset). The frequencies selected were between 8-13 Hz for alpha, and 15-25 Hz for beta (Coll et al., 2017). This additional analysis was done in order to match the univariate desynchronization data exactly to the frequency bands selected for the MVPA (see Multivariate pattern analysis below for comparison to MVPA). To test whether the ERSP data showed significant synchronization/desynchronization, we performed one-sample parametric *t*-tests against zero (baseline). All paired *t*-tests are reported as two-tailed at the *p* < .05 level with Bonferroni corrections applied. Effect sizes for all *t*-tests are calculated as Cohen’s *d* = *t* / √N.

#### Correlation analysis

In order to examine whether participants subjective ratings of sleepiness correlated with occipital alpha activity (Niedermeyer and da Silva, 2005), a correlation analysis was ran using participants scores from the KSS (see Apparatus and materials above) against all conditions and ROIs.

#### Multivariate pattern analysis

The MVPA was trained on single-trial ERSP data using a linear support vector machine (LIBSVM 3.20 toolbox; C. Chang & Lin, 2011) and tested against the average pattern from each visual object category (see Smith & Smith, 2019). The pattern classifiers were trained to discriminate between object identity within our two conditions: familiar or unfamiliar visual object categories. For example, in the familiar object condition, the classifier was trained to discriminate between an apple and a wine glass. A k-fold cross validation approach was used to estimate this performance, whereby the model was built from k – 1 subsamples (70% of trials) and tested on the average of the remaining independent k subsample (30% of trials). Therefore, the classifier was trained with 168 single trials (84 trials for each visual object category; e.g. apples and wine glasses), and tested on decoding performance against the average of 72 trials (36 for each visual object category) in each condition. This was carried out in both the alpha (8-13 Hz) and beta (15-25 Hz) frequency bands (see Coll et al., 2017; Cuellar & Toro, 2017), in both the central and occipital ROI’s, to test whether discriminable patterns of information could be detected for different familiar or unfamiliar visual objects in each ROI and frequency band. This analysis was performed on 20 randomly partitioned training test set iterations across the entire 1000ms stimulus duration time window. Overall, one decoding accuracy was obtained for each condition, separated by ROI and frequency band for each participant.

To test whether group level decoding accuracy was significantly above chance, we performed one-sample parametric *t*-tests on all MVPA analyses, against the chance level of 1/2 (50%). Significance values are reported as one-tailed due to prior expectations of the direction of the results. To control for multiple comparisons, all decoding accuracies are corrected using false discovery rate (FDR). The adjusted q-value at ≤ .05 resulted in a significance value of FDR *p* ≤ .016 for all results (Benjamini and Yekutieli, 2001). Effect sizes for all *t*-tests are calculated as Cohen’s *d* = *t* / √N. All effect sizes are to be identified as small (> .2), medium (> .5), and large (> .8) according to Cohen’s (1988) classification of effect sizes.

## Results

### Behavioural accuracy

The mean accuracy of catch trial detection was 99.56% (*SD* = 1.01%, range = 96% - 100%), thus indicating that participants were very good at detecting the absence or presence of a red and green fixation cross during a block. Participants sleepiness ratings covered the full scale, with an average of 5.74 (*SD* = 2.03, range 1 - 9). Average response times for the voluntary motor response task were 6838ms (*SD* = 933ms, range 5497 – 7244ms).

### Univariate results: Cluster-based analysis

For the univariate analysis, significant clusters were first identified via a data-driven approach, to investigate where there were significant differences in power across the 1-30 Hz frequencies. The purpose of this analysis was to determine whether a mu rhythm desynchronization could be detected from merely viewing the familiar graspable objects. We also tested whether a mu rhythm desynchronization could be detected in the voluntary motor response task.

#### Main experiment central ROI

When examining the significant clusters of both conditions averaged together in the central ROI, we observed significant synchronization over the delta/theta band covering stimulus duration and peaking around 100-400ms post stimulus onset, and significant desynchronization covering the beta band, also spanning stimulus duration and peaking over 200-600ms (see Figure 2A). Data from these clusters were then extracted separately for each condition (familiar or unfamiliar object categories; see Figure 2A). Note that the actual significant pixels for familiar and unfamiliar visual objects are displayed in Supplementary Figure 1, however to keep the number of pixels matched across both conditions and avoid circular inference, the masks based on the average of both conditions were used to extract the data (Figure 2A). The synchronization over the delta/theta band was found to be strongly significant for both familiar (*M* = .454, *t*_26_ = 7.918, *p* < .001, *d* = 1.524) and unfamiliar (*M* = .463, *t*_26_ = 6.174, *p* < .001, *d* = 1.188) visual object categories. Further pairwise comparisons revealed these were not significantly different from one another (*t*_26_ = .157, *p* = .876). The desynchronization from the beta band was also strongly significant for both familiar (*M* = −.272, *t*_26_ = −5.333, *p* < .001, *d* = −1.026) and unfamiliar (*M* = −.309, *t*_26_ = −6.252, *p* < .001, *d* = −1.203) visual object categories. Once again, further pairwise comparisons revealed these were not significantly different from one another (*t*_26_ = −1.207, *p* = .238). A bar chart displaying the averaged ERSP values can be seen in Figure 2B. Taken together, the results from the central ROI show that observation of graspable objects, regardless of familiarity with the object, causes desynchronization in the beta frequency band. In contrast to our expectations, there were no such significant effects found within the mu rhythm in these analyses.

**Figure 2:**
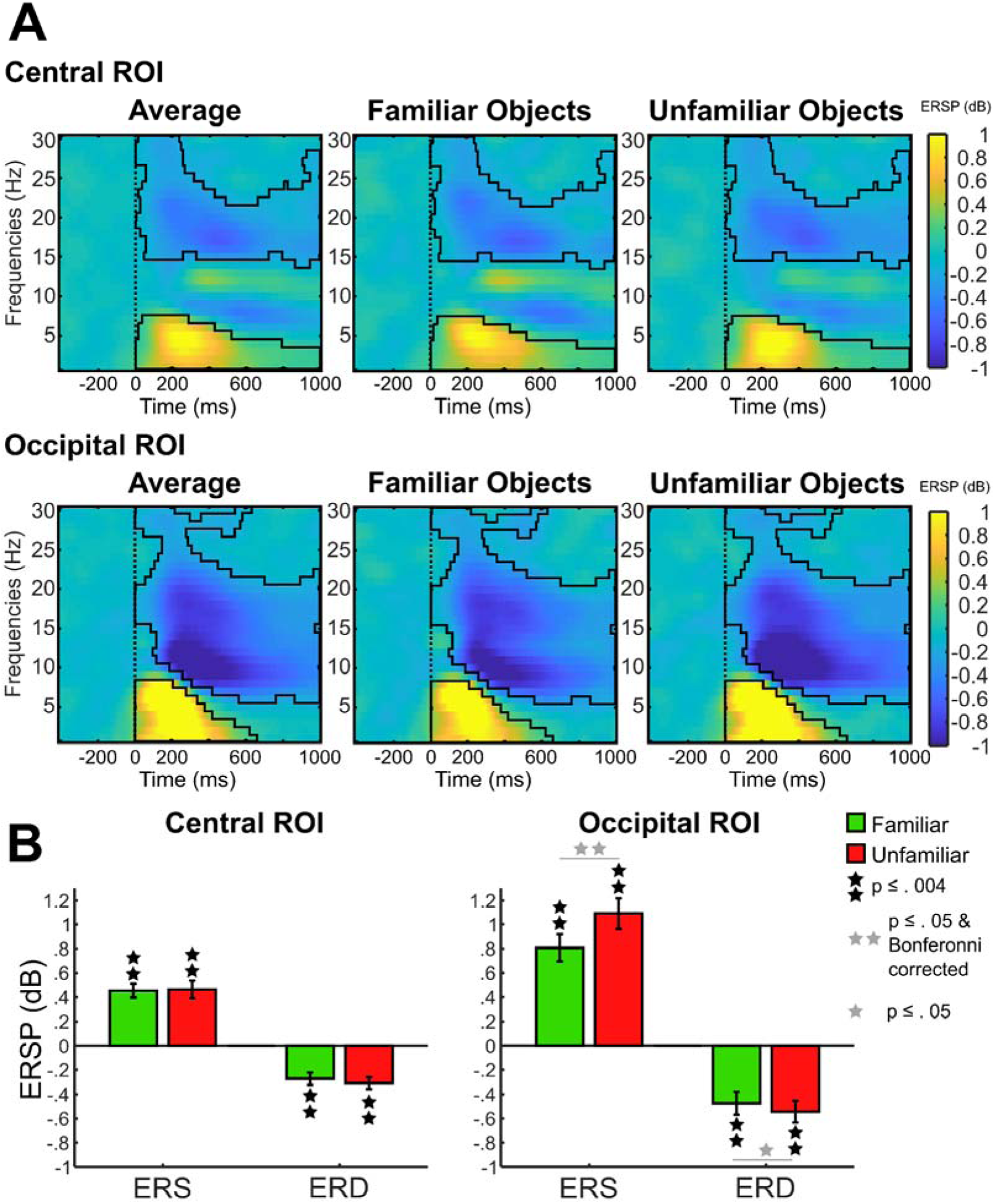
Univariate results from the cluster analysis. (A) The significant pixels taken from the average ERSP data from both familiar and unfamiliar objects, in both central and occipital ROIs. The raw ERSP data taken from this mask is visually shown for both conditions of familiar and unfamiliar visual objects. (B) Average ERSP values in each significant cluster for both central and occipital ROIs. Significant differences between conditions are shown.

#### Main experiment occipital ROI

When we examined the significant clusters of both conditions averaged together in the occipital ROI, we observed significant synchronization over the delta/theta band from 0-600ms and peaking between 0-400ms post stimulus onset, and significant desynchronization covering the alpha/beta band, spanning stimulus duration and peaking between 200-600ms (see Figure 2A). Data from these clusters were then extracted separately for each condition (familiar or unfamiliar object categories; see Figure 2A). Note that the actual significant pixels for familiar and unfamiliar visual objects are displayed in Supplementary Figure 1, however to keep the number of pixels matched across both conditions and avoid circular inference, the masks based on the average of both conditions were used to extract the data (Figure 2A). The significant synchronization covering the delta/ theta combined frequency bands was significant for both familiar (*M* = .807, *t*_26_ = 7.230, *p* < .001, *d* = .1.391) and unfamiliar (*M* = 1.090, *t*_26_ = 8.547, *p* < .001, *d* = 1.645) visual object categories. Here, mean synchronization for familiar and unfamiliar visual object categories were significantly different from one another (*t*_26_ = 3.298, *p* = .003, *d* = .635). The significant desynchronization covering the alpha and beta combined frequency bands was significant for familiar (*M* = − .474, *t*_26_ = −4.985, *p* < .001, *d* = −.959) and unfamiliar (*M* = −.543, *t*_26_ = −6.057, *p* < .001, *d* = −1.166) visual object categories. Further pairwise comparisons revealed the mean desynchronization for familiar and unfamiliar visual object categories were different from one another (*t*_26_ = −2.177, *p* = .039, *d* = −.419), however, this result did not survive Bonferroni correction at a corrected *p* value of .013. A bar chart displaying the exact averaged ERSP values can be seen in Figure 2B. The results in the occipital ROI suggest a strong desynchronization can be found over both the alpha and beta frequency bands in response to viewing both the familiar and the unfamiliar visual object categories, in which the suppression appears to be slightly stronger for viewing unfamiliar visual objects compared to familiar.

#### Voluntary motor response task central ROI

Three significant clusters were identified in the central ROI for the voluntary motor response task, as seen in Figure 3A. A significant desynchronization was found over the alpha band central electrodes (the mu rhythm), beginning at the onset of button press and lasting approximately 1000ms, with the peak around 400-800ms (*M* = −.712, *t*_26_ = −5.297, *p* < .001, *d* = −1.019). Another significant desynchronization was found over the beta frequency band, spanning around −100ms to 200ms time-locked to the button press, with the peak around 0-200ms (*M* = −.457, *t*_26_ = −6.419, *p* < .001, *d* = −1.235). Finally, significant synchronization was found over the beta frequency band, spanning around 500-1000ms post-button press, peaking around 700-1000ms (*M* = .590, *t*_26_ = 4.517, *p* < .001, *d* = .869). These results show a strong mu rhythm desynchronization can be found when participants completed a simple button-press experiment, demonstrating the mu rhythm can easily be detected when participants execute a physical motor response.

**Figure 3:**
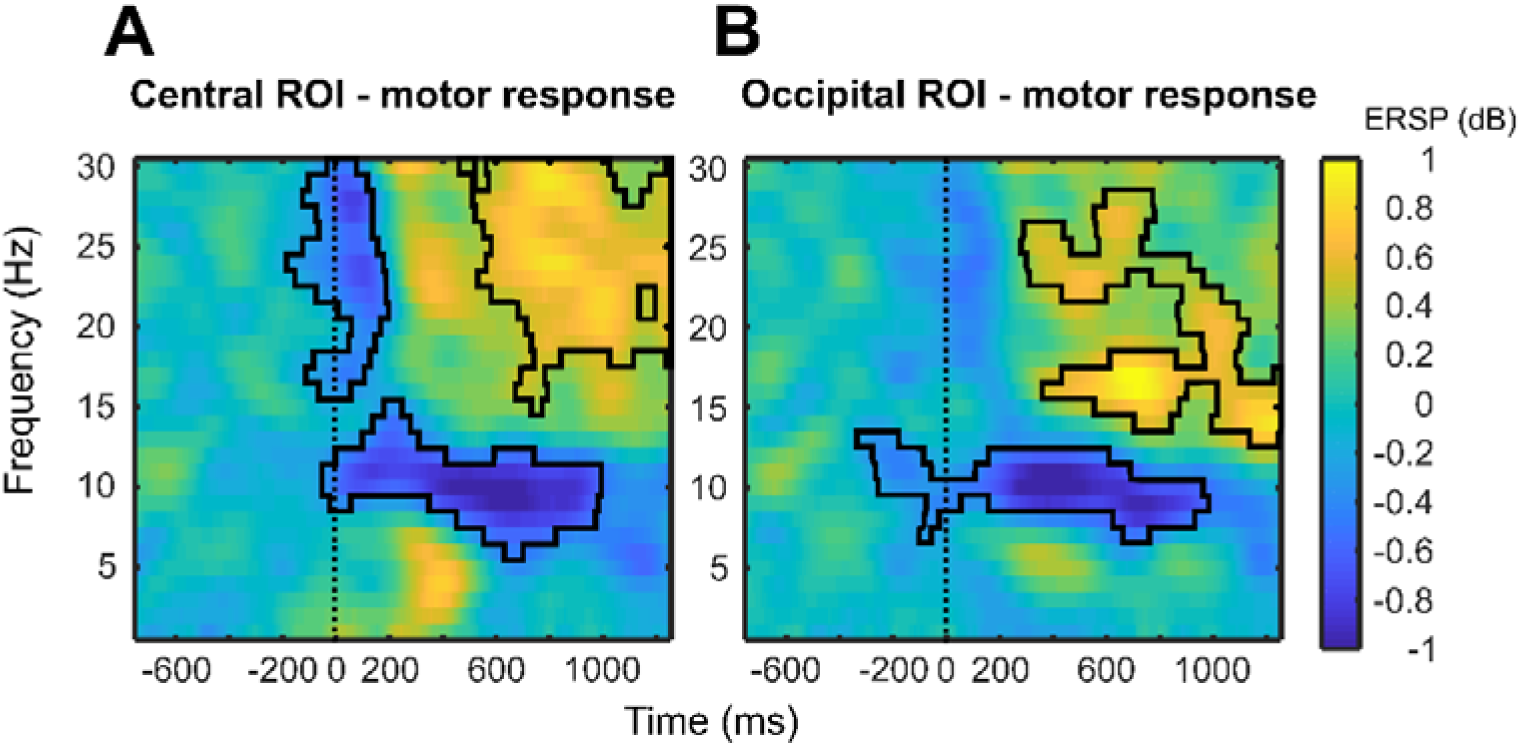
Univariate results from the cluster analysis. The significant pixels corresponding to the ERSP data from the voluntary motor response task are outlined in both central and occipital ROIs.

#### Voluntary motor response task occipital ROI

Two significant clusters were revealed in the occipital ROI for the voluntary motor response task (see Figure 3B). Significant desynchronization was found in the alpha frequency band from around −200ms to 1000ms time-locked to button press, and peaking around 200-600ms (*M* = −.653, *t*_26_ = −4.502, *p* < .001, *d* = −.866). Significant synchronization was also found in the beta frequency band, lasting from around 300-1000ms post-button press, and peaking around 600-800ms (*M* = .572, *t*_26_ = 4.469, *p* < .001, *d* = .860). These results show that the alpha band also attenuates in the occipital ROI in response to execution of a button press.

### Univariate results: Alpha- and beta-band analysis

The univariate data was also analysed within strictly selected alpha and beta frequency bands for each ROI (central and occipital) averaged over stimulus duration (0-1000ms time-locked to stimulus onset), in order to match the univariate desynchronization data exactly to the frequency bands selected for the MVPA.

#### Main experiment central ROI

When restricted to the 8-13 Hz frequency band, no significant desynchronization was found for either familiar (*t*_26_ = −.042, *p* = .967) or unfamiliar (*t*_26_ = −1.160, *p* = .257) visual object categories. Similar to the cluster-based analysis, the beta (15-25 Hz) frequency band revealed significant desynchronization for both the familiar (*M* = −.251, *t*_26_ = −4.630, *p* < .001, *d* = −.891) and unfamiliar (*M* = −.284, *t*_26_ = −5.296, *p* < .001, *d* = −1.019) visual object categories. Further pairwise comparisons revealed there were not significantly different from one another within the beta band (*t*_26_ = 1.001, *p* = .326). These results (see Figure 4) compliment the cluster-based analysis, suggesting no significant desynchronization within the mu rhythm frequency band, yet significant desynchronization for both visual object categories in the beta frequency band.

**Figure 4:**
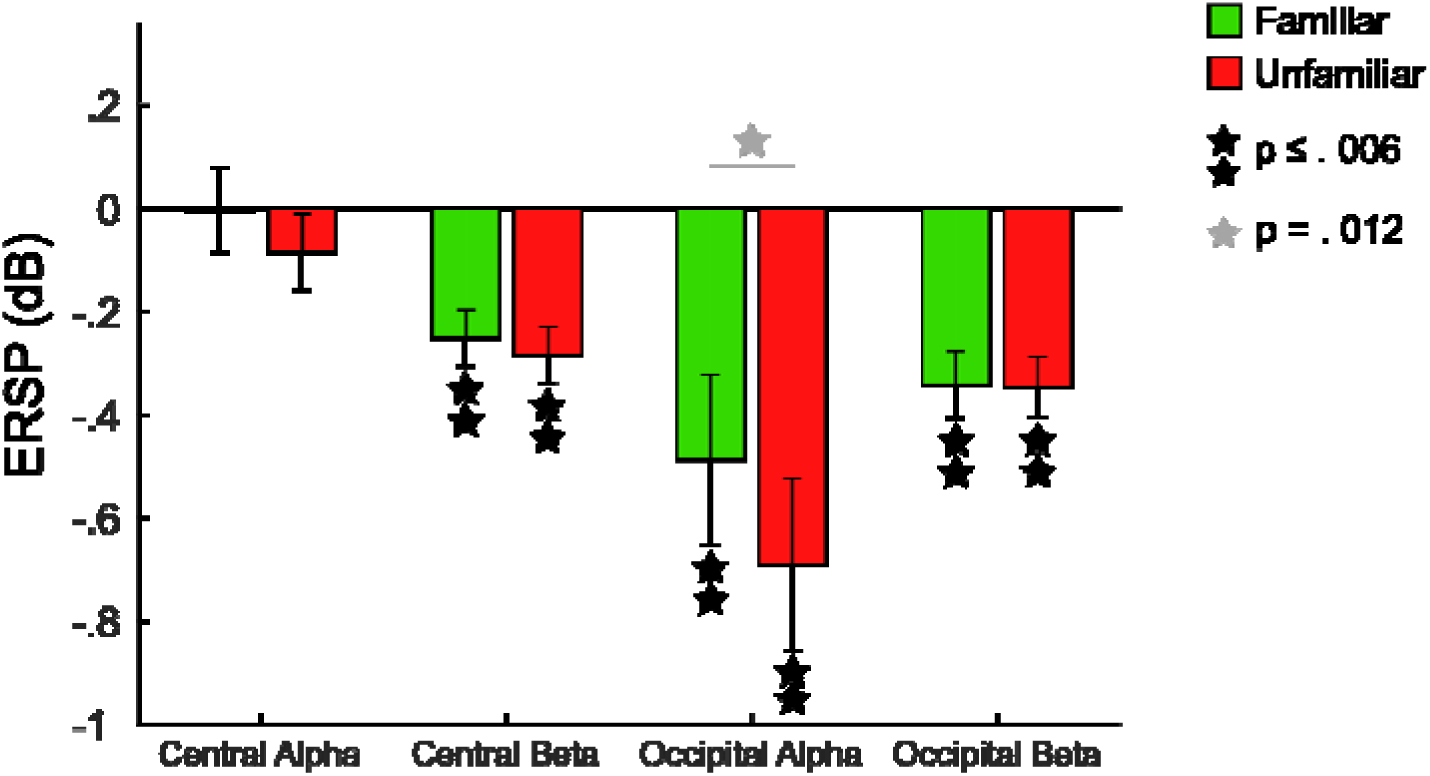
Univariate results from the alpha- and beta-band analysis. Figure shows the ERSP data in response to viewing both familiar and unfamiliar objects, in both central and occipital ROIs.

#### Main experiment occipital ROI

The 8-13 Hz frequency band in the occipital ROI revealed significant desynchronization for familiar (*M* = −.488, *t*_26_ = −2.972, *p* = .006, *d* = −.572) and unfamiliar (*M* = −.691, *t*_26_ = −4.161, *p* < .001, *d* = .801) visual object categories. Further pairwise comparisons revealed these to be significantly different from one another (*t*_26_ = 2.711, *p* = .012, *d* = .522). In the beta (15-25 Hz) frequency band, significant desynchronization was also found for both familiar (*M* = −.342, *t*_26_ = −5.336, *p* < .001, *d* = −1.027) and unfamiliar (*M* = −.346, *t*_26_ = −5.949, *p* < .001, *d* = −1.145) visual object categories. Further pairwise comparisons revealed these were not significantly different from one another (*t*_26_ = .164, *p* = .871). These results (see Figure 4) compliment the cluster-based analysis, showing alpha and beta desynchronization for both visual object categories. Interestingly, when separating the alpha and beta band rather than looking at the combined cluster, we find the alpha band shows significant differences in desynchronization for the familiar and unfamiliar visual objects, with stronger attenuation for viewing the unfamiliar objects compared to the familiar objects.

### Correlation analysis

The results from the correlation analysis revealed no significant correlations between each participant’s subjective sleepiness ratings and any alpha activity across all conditions and ROI’s (all *p*’s ≥ .321). As such, we can assume participants feelings of drowsiness throughout the experiment did not influence any of the alpha desynchronizations that we observe.

### Multivariate pattern analysis results

In order to determine whether content-specific information regarding the familiar visual objects could be decoded from the mu rhythm frequency band, we computed cross-validated decoding performance of visual object category independently for each condition (familiar and unfamiliar visual objects) in the central and occipital ROI, for both the alpha (8-13 Hz) and beta (15-25 Hz) frequency bands. The analysis was conducted across stimulus duration (0-1000ms time-locked to stimulus onset). Therefore, this analysis matched the univariate alpha- and beta-band analysis (see Univariate results: Alpha- and beta-band analysis above).

#### Central ROI

Significantly above chance decoding was found for the familiar visual object categories in the central alpha (mu rhythm) frequency band (*M* = 57.31%, *t*_26_ = 2.268, *p* = .016, *d* = .436; see Figure 5). Conversely, such decoding effects were not found for the unfamiliar object categories (*t*_26_ = .670, *p* = .509). Further paired samples tests showed these decoding accuracies were not significantly different from one another (*t*_26_ = .921, *p* = .366). Interestingly, no significant decoding was found for either familiar (*t*_26_ = −.746, *p* = .462) or unfamiliar (*t*_26_ = .026, *p* = .980) visual object categories in the central beta frequency band. These results show, despite a lack of desynchronization in the mu rhythm in the univariate analysis, discriminable information regarding the familiar, but not unfamiliar, visual object categories can be found in the mu rhythm oscillatory response.

**Figure 5:**
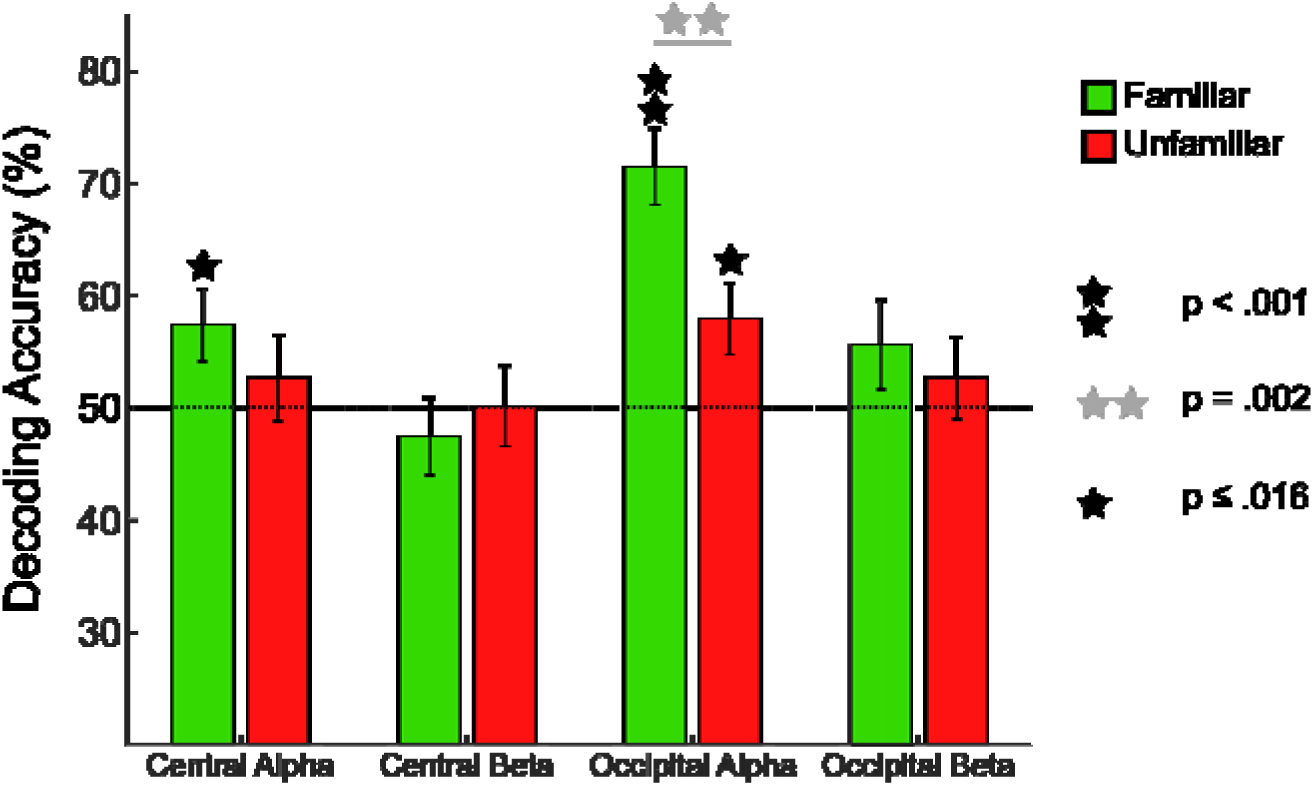
Decoding of object identity within each visual object category. Cross-validated two automatic forced choice decoding performance for each stimulus category (familiar and unfamiliar objects) for each frequency band and ROI.

#### Occipital ROI

As can be seen in Figure 5, strong significant decoding was found for familiar visual object categories in the occipital alpha frequency band (*M* = 71.48%, *t*_26_ = 6.284, *p* < .001, *d* = 1.209). Significant decoding was also found for unfamiliar object categories (*M* = 57.87%, *t*_26_ = 2.478, *p* = .010, *d* = .477). Interestingly, further comparisons revealed decoding for familiar visual object categories to be significantly higher than decoding for unfamiliar visual object categories (*t*_26_ = 3.124, *p* = .002, *d* = .601). No significant decoding was found for either familiar (*t*_26_ = 1.385, *p* = .178) or unfamiliar (*t*_26_ = .699, *p* = .491) visual object categories in the occipital beta frequency band. These results show discriminable information for both conditions of familiar and unfamiliar visual object categories in the alpha frequency band, which compliments the desynchronization results in the univariate analysis. However, interestingly the familiar visual objects are significantly more discriminable than the unfamiliar visual objects, which is in contrast to the univariate analysis which reveals stronger desynchronization for the unfamiliar objects in the occipital alpha frequency band. Decoding was not significant for either object category in the beta frequency band, which is interesting given strong significant desynchronization was found within the beta band in the univariate analysis.

## Discussion

The present study demonstrates, for the first time, that the mu rhythm can produce specific patterns of activity in response to viewing images of different familiar objects. This was found despite no tactile stimulation or actions being performed or contained in the stimuli. Interestingly, no reliable information related to viewing different images of unfamiliar graspable objects was found in the mu rhythm. Thus, viewing images of familiar graspable objects may automatically trigger activation of associated tactile or haptic properties in sensorimotor areas. Furthermore, univariate analysis investigating changes in event related spectral power revealed significant attenuation in the central beta frequency band when viewing both familiar and unfamiliar graspable objects, whilst no such attenuation was found in the mu rhythm oscillatory response. Finally, our analyses of occipital alpha showed an interesting pattern of greater attenuation for unfamiliar objects but better representations for familiar object categories, in line with certain account of how expectations influence neural processing (de Lange et al., 2018). On a methodological level our findings further highlight how the analysis technique employed (univariate vs multivariate) can offer complementary views on the role of neural oscillations in response to viewing different visual object categories.

### The mu rhythm contains information that permits discrimination of familiar graspable objects from vision

In the current experiment we show that information in the mu rhythm can discriminate different familiar object categories, but not unfamiliar categories. As such we build and extend upon prior work that has shown the mu rhythm contains information about observed and executed actions (Coll et al., 2017). Importantly in this prior study the authors classified features of observed and executed actions such as transitivity (object directed vs mimed), grasp type (whole hand vs precision) and the presence of tactile vibration (on vs off). Our work goes beyond this by showing that the mu rhythm contains information about object identity for familiar graspable objects – despite the absence of any observed or performed object actions in the experiment.

While prior work using univariate methods has found greater attenuation of the mu rhythm to pictures of tools vs non-tools (Proverbio, 2012) and to real 3D vs 2D pictures of objects (Marini et al., 2019), in the current experiment we did not find any difference in terms of attenuation between familiar and unfamiliar object categories. This may be because both our familiar and unfamiliar categories comprise pictures of graspable object categories. In addition, our familiar objects were limited in number (3 examples of 2 object categories) and did not include tool examples but rather everyday common objects. While there are advantages in using tools such as the action routines being highly stereotyped to these objects (e.g. Knights et al., 2022, 2021) here we favoured objects that had richer multisensory knowledge in the expectation that contextual mechanisms – underpinning transmission of information from visual to sensorimotor areas - are likely to work most strongly in this case (see Smith and Goodale, 2015; Smith and Muckli, 2010 for related arguments).

These findings also resonate strongly with our prior fMRI study that show that familiar but not unfamiliar objects can be discriminated within primary somatosensory brain areas (Smith and Goodale, 2015; see also Bailey et al., 2023). This study used a superset of the objects used in the current EEG experiment and revealed a comparable pattern of results – reliable decoding of the identity of familiar graspable objects but not unfamiliar graspable objects from primary somatosensory areas. Crucially our previous study also showed no reliable decoding effects in M1 for familiar or unfamiliar objects (Smith and Goodale 2015). Taken together with past work that has shown that the mu rhythm may be more related to tactile stimulation than action per se (Coll et al., 2017) and may localize to primary somatosensory cortex rather than motor cortex (Cheyne et al., 2003; Ritter et al., 2009) it is likely that we are measuring in both cases the same underlying effect and that the generators of our measured decoding effects on the mu rhythm are in SI. However, in the absence of source localization we acknowledge that this interpretation remains somewhat speculative. Nonetheless both studies provide strong converging evidence for the view that upon sight of familiar graspable objects information is triggered in early sensorimotor brain areas related to object identity. If predictive processing is taken to be the computational goal of the brain (Clark, 2013; Friston, 2009) then it may be the case that viewing familiar graspable objects (or in fact hearing sounds associated with such objects – Bailey et al., 2023) automatically leads to fine grained information about associated tactile or haptic features of such objects being activated in early somatosensory (or sensorimotor) areas (see Pérez-Bellido et al., 2018 for a related argument; see also Avery et al., 2021).

An alternative reason why we find information specific to the content of only familiar visual objects in the mu rhythm may be due to the fact participants paid more attention to an object they were familiar with. However, since spectral power changes in the occipital alpha oscillation are sensitive to attentional engagement (Hobson and Bishop, 2016), and we actually found stronger desynchronization for viewing the *unfamiliar* objects compared to the familiar objects, this suggests it is unlikely that we are merely measuring engagement of attention to objects that participants are more familiar with.

We emphasize the fact that the mu rhythm can be used to answer more questions beyond the function of the mirror neuron system (see also Proverbio 2012; Marini et al., 2019), and demonstrate how different analysis methods can provide complementary insights into the key function of the mu rhythm oscillatory response. For example, our research suggests the content of information which is likely being sent to sensorimotor cortex via feedback connections from a distal area of cortex (in our case, visual cortex or high-level multisensory convergence zones) can be measured in the mu rhythm when using MVPA techniques. This information may not be detected when using simpler univariate analysis such as assessing attenuation of the mu rhythm. This is important to note because most previous research has analysed mu rhythm attenuation to measure sensorimotor cortex activation. To directly compare this to our results would suggest that viewing and executing actions, or receiving tactile stimulation, is pivotal to detect a mu rhythm response. This is because no significant attenuation was found in the mu rhythm when viewing different images of either familiar or unfamiliar graspable objects in the present study, whereas we find strong significant attenuation when participants performed a voluntary motor response task. However, a crucial finding in our research is the fact that we did find discriminable information about the different familiar graspable objects in the mu rhythm when analysing the data at the multivariate level. This result highlights how such research should consider adopting a strong focus on multivariate classification techniques to further understand the role of the mu rhythm (see also Coll et al., 2017), since this method has higher sensitivity and power to detect fine-grained differences in the representational content of the data (see Norman, Polyn, Detre, & Haxby, 2006).

### The central beta oscillatory response to observation of graspable objects

The univariate analysis revealed significant desynchronization in the central beta oscillation in response to participants viewing both familiar and unfamiliar graspable objects. These responses were not significantly different from one another, suggesting there are no familiarity effects in the central beta band, but rather viewing any graspable object in general can elicit central beta band desynchronization. The significant attenuation is not a surprising finding given previous research has found the central beta oscillation is involved in action related processes such as motor imagery, passive movement, and action observation (Zaepffel et al., 2013). However, interestingly, whilst significant desynchronization was observed in the central beta band when viewing familiar and unfamiliar graspable objects, multivariate analysis could not reliably discriminate between the graspable objects for either category in this frequency band. Therefore, it seems the central beta oscillation can classify between an observed or executed action (Coll et al., 2017), and may even play a role in movement planning (see also Tucciarelli, Turella, Oosterhof, Weisz, & Lingnau, 2015; Turella et al., 2016), however it fails to distinguish between different action types or different tactile properties of objects. Here, these results once again highlight the importance of using different analysis techniques to answer different questions about oscillatory responses.

Another interesting finding is the clear difference of attenuation found between the main experiment and the voluntary motor response task in the central beta oscillation. Neuper, Wörtz, and Pfurtscheller (2006) found beta band suppression during preparation of movement, followed by a strong rebound beta synchronisation after movement, which occurs whilst the mu rhythm continues to desynchronise. This is exactly what we see in the voluntary motor response task and is important to highlight for two reasons. First, this directly indicates how central alpha and beta frequency bands exhibit different dynamics (as previously suggested by Pfurtscheller et al 1994), emphasizing the importance of separating these frequency bands to answer different questions regarding cortical function. This contrasts with previous research which suggests the central alpha (mu rhythm) and central beta frequency bands are closely related to one another during action production and gesture observation (Quandt et al., 2012). Future research should consider an EEG source-based analysis or magnetoencephalography (MEG) study in order to estimate the location of neural activation found in the alpha and beta frequency bands. Indeed, previous research has found mu and beta correspond to different sources in SI and MI (Cheyne, 2013; Cheyne et al., 2003; Hari and Salmelin, 1997). Based on our results, one might argue that the mu rhythm desynchronization corresponds to the tactile feel of the button press in SI, whereas the beta desynchronization corresponds to the motor plan of the button press in MI or pre-motor cortex (Tucciarelli et al., 2015; Turella et al., 2016). Second, the difference in desynchronization between the main experiment and the voluntary motor response task emphasizes the importance of separating observation and execution conditions to answer different questions about the central beta oscillation, in line with Coll et al. (2017) who suggested the central beta oscillation can classify between an observed or executed action, yet fails to show specificity to different action types, such as an action with a real object or pantomime action.

### Occipital alpha may reflect top-down synchronous activity when viewing graspable objects

The results found in the occipital alpha frequency band may reflect top-down neuronal processes underlying perception of objects, since the alpha frequency has previously been suggested to play an important role in directing information flow through the brain and allocating resources to relevant brain regions (Haegens et al., 2011; Jensen and Mazaheri, 2010). Our univariate analysis revealed significant desynchronization in the occipital alpha/beta cluster in response to viewing images of both familiar and unfamiliar graspable objects, in which the suppression was significantly stronger for unfamiliar objects in the alpha band when restricting the analyses to precise frequency boundaries. The overall attenuation may be a reflection of increased blood flow to the visual cortex during perception, since Perry and Bentin (2009) have previously found a relationship between alpha suppression recorded posteriorly and BOLD responses in parietal and visual cortex. Scheeringa et al. (2011) also found alpha suppression reflects the magnitude of the MRI response in visual cortex during a visual attention task. The reason why attenuation was stronger for the unfamiliar objects in the occipital alpha band may be a result of a novelty effect, since previous research has demonstrated stronger occipital alpha desynchronization following presentation of a novel stimulus compared to a familiar expected stimulus (Harrison, 1946; Mulholland and Runnals, 1962).

We argue this difference is not simply a confound of expectation about the onset of the stimulus or attentional engagement, which are both known to influence alpha activity (see e.g. Hobson & Bishop, 2016; Pfurtscheller, 1992). Tight controls were adopted in this experiment to account for such confounds; for instance, maintaining constant fixation with a variable delay eliminated the risk of forming an expectation about when the stimuli might appear (Samaha et al., 2015). Furthermore, the results of the correlations between KSS scores (Akerstedt et al., 2014; Åkerstedt and Gillberg, 1990) and occipital alpha desynchronization suggest no significant relationship between feelings of sleepiness, therefore arousal, and the power of attenuation in the occipital alpha frequency band. Finally, if the stronger suppression in the occipital alpha oscillation for viewing unfamiliar objects compared to familiar objects is due to stronger attention paid to these stimuli, we may expect to find a significantly stronger suppression in the mu rhythm also. This is because Hobson and Bishop (2016) argue the mu rhythm is easily confounded with occipital alpha suppression. However, we do not find the same pattern of results in the mu rhythm oscillatory response, with the strength of desynchronization between viewing familiar and unfamiliar objects not being significantly different from one another.

Interestingly, the multivariate analysis in the occipital alpha band could significantly discriminate within both the familiar and unfamiliar visual object categories, whereby decoding is significantly stronger for viewing *familiar* graspable objects when compared to unfamiliar. We suggest this difference is due to the two analysis techniques detecting different aspects of the oscillatory response, since multivariate analysis has more power to detect fine-grained differences in the *representational* content of the data (Norman et al., 2006). As mentioned previously, Bastos et al. (2012) suggested the alpha frequency coordinates the feedback of predictions to low-level areas, however occipital alpha in particular is known to transmit prior knowledge and expectations to visual cortex, such as in a perceptual decision making experiment (Sherman et al., 2016; see also Chen et al., 2023). As such, it may be the case that familiar objects could be better read out from occipital alpha activity due to the availability of previous knowledge for these objects. Intriguingly the pattern of effects found on alpha attenuation vs representation are in keeping with the sharpening account of expectation suppression (de Lange et al., 2018; Kok et al., 2012), implying that familiarity with the object leads to greater attenuation but better representation.

It is important to note that the direction of the multivariate effect in the occipital alpha band is similar to the multivariate effect in the central alpha band (mu rhythm), meaning in both cases decoding is stronger for viewing familiar compared to unfamiliar objects. The reason why the effect is similar (yet weaker) in the mu rhythm may be a case of more information being fed back to occipital cortex than sensorimotor cortex regarding the stored knowledge about the familiar *visual* objects (Martin, 2016), since feedback information is suggested to oscillate at an alpha frequency (Bastos et al., 2012). To confirm this idea, an interesting avenue for future research could consider conducting the multivariate analyses in the gamma (40-80 Hz; Seymour, Rippon, Gooding-Williams, Schoffelen, & Kessler, 2018) frequency range in both occipital and central electrodes, since information processing in this frequency is coherent with activity in superficial layers of cortex (Buffalo et al., 2011). This is important since feedforward connections predominantly arise from superficial layers of cortex (Barone et al., 2000; Buffalo et al., 2011). Furthermore, research has suggested gamma conveys feedforward information (Van Kerkoerle et al., 2014). Therefore, if gamma reflects feedforward processing, we would expect to find above chance decoding for both familiar and unfamiliar graspable objects in the occipital gamma oscillation, yet no above chance decoding in the central gamma oscillation. Furthermore, we would expect to find no differences between decoding accuracies for familiar and unfamiliar graspable objects in the occipital gamma band if this frequency reflects feedforward input, since Smith and Goodale (2015) found no significant differences in decoding between viewing familiar and unfamiliar graspable objects in VI. This idea is further supported by previous research which has found gamma activity most strongly and significantly contributed to explaining BOLD variance (Scheeringa et al., 2011) or changes in contrast (Koch et al., 2009) in human visual cortex.

## Conclusion

In summary, this study has shown that the mu rhythm oscillation contains information specific to the identity of familiar object categories, when triggered via vision. In contrast, no such effects were found when participants viewed images of unfamiliar graspable objects. Our work provides converging evidence that the neural representations present in early sensorimotor areas (most likely primary somatosensory cortex) can be shaped in a fine grained manner by relevant contextual information originating in different sensory modalities. As such we have shown, for the first time, evidence for the precise temporal communication of information, and thus a potential oscillatory marker, of cross-sensory effects from vision to sensorimotor cortex. Thus our approach offers a novel way to quantify the neural influences of object affordances in future research. Finally, we highlight the importance of using different analysis techniques to extract different types of information from neural oscillations. We emphasize the need for research to focus on multivariate analysis techniques which can read out fine-grained pattern information from oscillatory responses, in turn answering new questions that simple analyses on attenuation of power fail to detect.

---

**Supplementary Figure 1:**
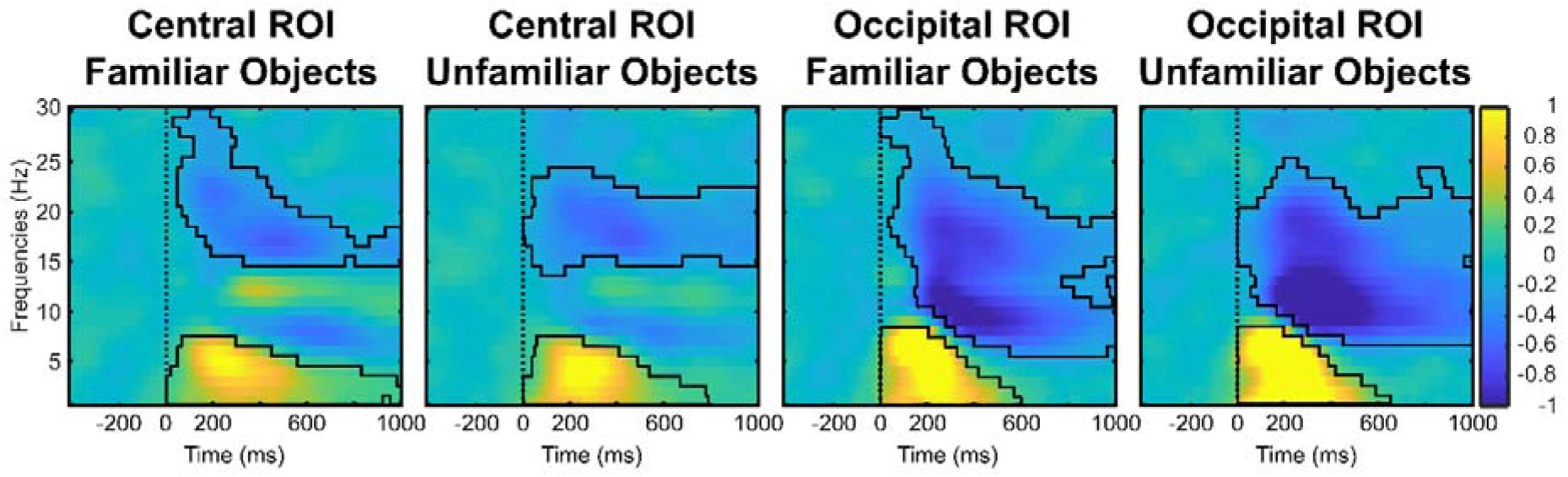
Masks of the actual significant pixels from the cluster-based analysis for each visual object category condition and ROI.

